# *RAB23* loss-of-function mutation causes context-dependent ciliopathy in Carpenter syndrome

**DOI:** 10.1101/2025.02.10.637381

**Authors:** WY Leong, WL Tung, WH Chui, Andrew O M Wilkie, CHH Hor

## Abstract

The primary cilium is a signal transduction organelle whose dysfunction clinically causes ciliopathies in humans. RAB23 is a small GTPase known to regulate the Hedgehog signalling pathway and ciliary trafficking. Mutations of *RAB23* in humans lead to Carpenter syndrome (CS), an autosomal recessive disorder clinically characterized by craniosynostosis, polysyndactyly, skeletal defects, obesity, and intellectual disability. Although the clinical features of CS bear some resemblance to those of ciliopathies, the exact relationship between the pathological manifestations of CS and the ciliary function of *RAB23* remains ambiguous. Besides, the *in vivo* ciliary functions of *RAB23* remain poorly characterised.

Here, we demonstrate *in vivo* and *in vitro Rab23* loss-of-function mutants modelling CS, including *Rab23* conditional knockout (CKO) mouse mutants, CS patient-derived induced pluripotent stem cells (iPSCs), and zebrafish morphants. The *Rab23*-CKO mutants exhibit multiple developmental and phenotypical traits recapitulating the clinical features of human ciliopathies and CS, indicating a causal link between the loss of *Rab23* and ciliopathy. In line with the ciliopathy-like phenotypes, all three different vertebrate mutant models consistently show a perturbation of primary cilia formation, intriguingly, in a context-dependent manner. *In vivo* examination of primary cilia in *Rab23*-CKO mutants reveals profound cell-type specific ciliary abnormalities in chondrocytes and neocortical neurons, but not in epithelial cells, cerebellar granule cells and hippocampus neurons. A profound reduction in ciliation frequency and/or shortening of primary cilia was observed in the neurons and neural progenitor cells derived from CS patient iPSCs. Furthermore, *Rab23*-KO neural progenitor cells were desensitized to primary cilium-dependent activation of the Hedgehog signaling pathway. Collectively, these findings indicate that the absence of *RAB23* causes dysfunctional primary cilia in a cell-type distinctive manner, which underlies the pathological manifestations of CS. Our findings present the first *in vivo* evidence validating the unique context-specific function of *RAB23* in the primary cilium. Through the use of patient-derived iPSCs differentiated cells, we present direct evidence of primary cilia anomalies in CS, thereby confirming CS as a ciliopathy disorder.

## Introduction

The primary cilium is a microtubule-based elongated protrusion found on the surface of nearly all quiescent or differentiated mammalian cells. The primary cilium axoneme harbors a myriad of sensory and transmembrane receptors, making it a crucial signal transduction hub in cellular physiology (Gerdes et al., 2009; Silva & Cavadas, 2023; Wheway et al., 2018). In humans, dysfunction of the primary cilium results in a range of rare multisystemic congenital disorders that are collectively known as ciliopathies (Reiter & Leroux, 2017). Notably, ciliopathy disorders often exhibit overlapping phenotypes albeit genetically distinct. For example, Bardet–Biedl (BBS) and Joubert (JBTS) syndromes are ciliopathies caused by different gene mutations but share multiple common phenotypes such as retinal degeneration, cerebellar malformation and cognitive impairment, polycystic kidneys, and polydactyly (Waters & Beales, 2011).

Carpenter syndrome (CS) is a pleiotropic autosomal recessive genetic disorder often reported in patients carrying biallelic pathogenic variants in *RAB23* (Alessandri et al., 2010; Haye et al., 2014; Jenkins et al., 2011). The disorder is typically characterized by craniosynostosis, finger and toe deformity (such as polydactyly, brachydactyly, and syndactyly), visual and hearing impairments, short stature, obesity, and mild to severe intellectual disability. Other variable clinical presentations observed in CS patients include heart defects, molar agenesis, hypogenitalism, genu valgum and hydrocephaly (Hidestrand et al., 2009; Jenkins et al., 2007, 2011; Kadakia et al., 2014; Lodhia et al., 2021; Tarhan et al., 2004). Notably, numerous common clinical features of CS patients are overlapping with the common clinical hallmarks of ciliopathies, such as polysyndactyly, hydrocephaly, obesity, and intellectual disability. Despite these clinical similarities and recent speculations on the association between *RAB23* and ciliopathy (Zhao et al., 2023), it remains unclear whether patients with CS exhibit defective primary cilia.

Rab23 is a member of the small GTPase family, that forms a GDP-GTP exchange factor (GEF) complex with planar cell polarity effectors, Inturned and Fuzzy (Gerondopoulos et al., 2019). In mouse null mutant models, Rab23 was found to negatively regulate the Sonic hedgehog (Shh) signaling pathway (Eggenschwiler et al., 2001). The Shh signaling pathway is predominantly regulated in a primary cilium-dependent manner. In this process, the presence of Shh ligands induces Smoothened (Smo) to translocate to the primary cilium axoneme, which subsequently initiates the downstream signaling cascade through Gli transcription factors (Denef et al., 2000). Although Rab23 has been shown to mediate the ciliary localisation of Smo, Dopamine receptor 1 (D1R), and Kif17 (Boehlke et al., 2010; C. H. H. Hor et al., 2021; Leaf & Von Zastrow, 2015; Lim & Tang, 2015), the roles of *Rab23* in primary cilia biogenesis and/or maintenance remain ambiguous and controversial. Independent groups have reported inconsistent results on the effect of siRNA-directed silencing of *Rab23* in mouse inner medullary collecting duct (IMCD3) cells. While Gerondopoulos et. al observed a reduced degree of ciliation, Leaf and von Zastrow reported an unperturbed extent of ciliation in *Rab23-*knockdown IMCD3 cells (Gerondopoulos et al., 2019; Leaf & Von Zastrow, 2015). A similar discrepancy was also observed in the immortalized retinal pigmented epithelial (hTERT-RPE1) cell line. Two independent groups have reported a reduced ciliation (Gerondopoulos et al., 2019; Yoshimura et al., 2007), whereas another study observed no discernible change in the percentage of ciliation, despite using the same siRNA-mediated knockdown approach to silence *Rab23* in hTERT-RPE1 cells (Lim & Tang, 2015). However, in our previous study on *Rab23-*knockout cerebellar primary granule progenitor cells culture (C. H. H. Hor et al., 2021), we observed a reduction in the length of primary cilia, albeit with an unperturbed rate of ciliation. Given these inconsistent observations on the effect of *Rab23* silencing on the primary cilia formation and its structural integrity, more extensive investigations are necessary to elucidate the precise functions of Rab23 within the primary cilium.

In this study, we have established novel disease models of CS using transgenic mice and human induced pluripotent stem cells (iPSCs) reprogrammed from biopsies of CS patients. Strikingly, both the global gene knockout (KO) mutants (actin-Cre), and neural progenitor cells (npc)-specific knockout mutants of *Rab23* display a range of developmental and phenotypic abnormalities that largely resemble the clinical features observed in CS and ciliopathy patients. In these mutants, a significantly reduced number of primary cilia were observed in the cerebral neocortical neurons, suggesting that Rab23 plays important ciliary functions in the neocortical neurons. Intriguingly, the frequency of ciliated cells appeared largely unchanged in other brain regions such as the hippocampus neurons and granule precursor cells in the cerebellum, or in other non-neuronal cell types such as the epithelial cells and chondrocytes; however, *Rab23-*KO neural progenitor cells and chondrocytes show altered cilia length and volumes. Similar observations were found in *rab23*-silenced zebrafish morphant (*rab23* MO), in which the frequency of ciliation in the neurons was predominantly perturbed in the rostral neural tube but not in other regions such as the more caudally located spinal cord neural tube nor the pronephric duct. Consistent with the evidence from mouse and zebrafish mutant models, primary cilia anomalies were also observed in human neural progenitor cells and neurons differentiated from a CS patient’s iPSCs.

Collectively, our findings demonstrate that primary cilium anomalies occur more predominantly in the *RAB23* loss-of-function neurons (albeit in distinct neuronal populations) and this phenomenon is consistently observed in the zebrafish morphants, transgenic KO mice, and human disease models of CS patients, suggesting that the function of *RAB23* in the primary cilium is context-dependent and evolutionarily conserved. In summary, the collective results from various animal and cellular disease models indicate that tissue- and cell-type-specific dysfunctions of primary cilia could contribute to the pathological features of Carpenter syndrome, implying that Carpenter syndrome is a ciliopathy disorder. Moreover, *RAB23* appears to play crucial roles in the maintenance and/or biogenesis of primary cilia.

## Results

### RAB23 is highly conserved across different vertebrates

In order to examine if RAB23 is conserved amongst vertebrates, we performed multiple amino acid sequence alignment to examine the similarity of the *RAB23* encoding region across different vertebrates (Supplementary Figure 1). Our data showed that the amino acid sequences in human, macaque (*Macaca mulatta*), mouse (*Mus musculus*) and zebrafish (*Danio rerio*) display high degrees of similarity, with 93.25 % sequence identity between human and mouse, and 84.39 % between human and zebrafish respectively (Supplementary Figure 1). These results indicate that the RAB23 sequence is highly conserved across vertebrates. Given the high similarity in the amino acid sequences of RAB23 from humans to zebrafish, we generated zebrafish morphants and mouse mutants of *Rab23* to enable elucidation of Rab23’s role in the functional integrity of the primary cilium and the consequence of its loss in the pathogenesis of Carpenter syndrome.

### *Rab23*-KO mouse disease models of Carpenter syndrome phenocopies ciliopathy

In order to generate transgenic knockout mouse mutants of *Rab23*, *Rab23^f/f^* homozygous mice (C. H. H. Hor et al., 2021) were crossed with a β-actin-Cre driver line to induce global deletion of *Rab23* in the progenies (herein named actin-CKO). The first reported null mutant of *Rab23,* i.e. the ENU-induced *opb2* mouse mutant exhibited embryonic lethality at embryonic day (E) 12.5 (Eggenschwiler et al., 2001). Recent work has also reported an *opb2* mutant with an extended prenatal survival to E18.5 by breeding the mouse line into a C57BL6 strain (Hasan et al., 2020). Similar to the latter *opb2* mutant, our actin-CKO mutant was viable throughout the prenatal stage in the C57/BL6 background but died postnatally. The actin-CKO embryos at E12.5 exhibited polysyndactyly, missing or abnormal eyes and morphologically aberrant posterior neural tube (Fig. 1A). Furthermore, the actin-CKO embryo at E18.5 exhibited several prominent phenotypic characteristics that closely resembled those observed in individuals diagnosed with human Carpenter syndrome (Alessandri et al., 2010; Haye et al., 2014; Jenkins et al., 2007, 2011). These characteristics encompass craniofacial anomalies, polysyndactyly, growth retardation, and brain deformities (Fig. 1A-D). Notably, a range of abnormal brain morphologies, varying from mild to severe (data not shown), were observed. The milder cases displayed thinning and mis-patterning of the cerebral cortex (Fig. 1B-D), while an altered pattern of the cerebellar anlage at E18.5 was also evident (Fig. 1B, indicated by red asterisks). Importantly, these abnormalities largely align with the common phenotypic features found in previously reported ciliopathy mutants. (Damerla et al., 2015; Oud et al., 2016; Youn & Han, 2018).

**Figure 1.**
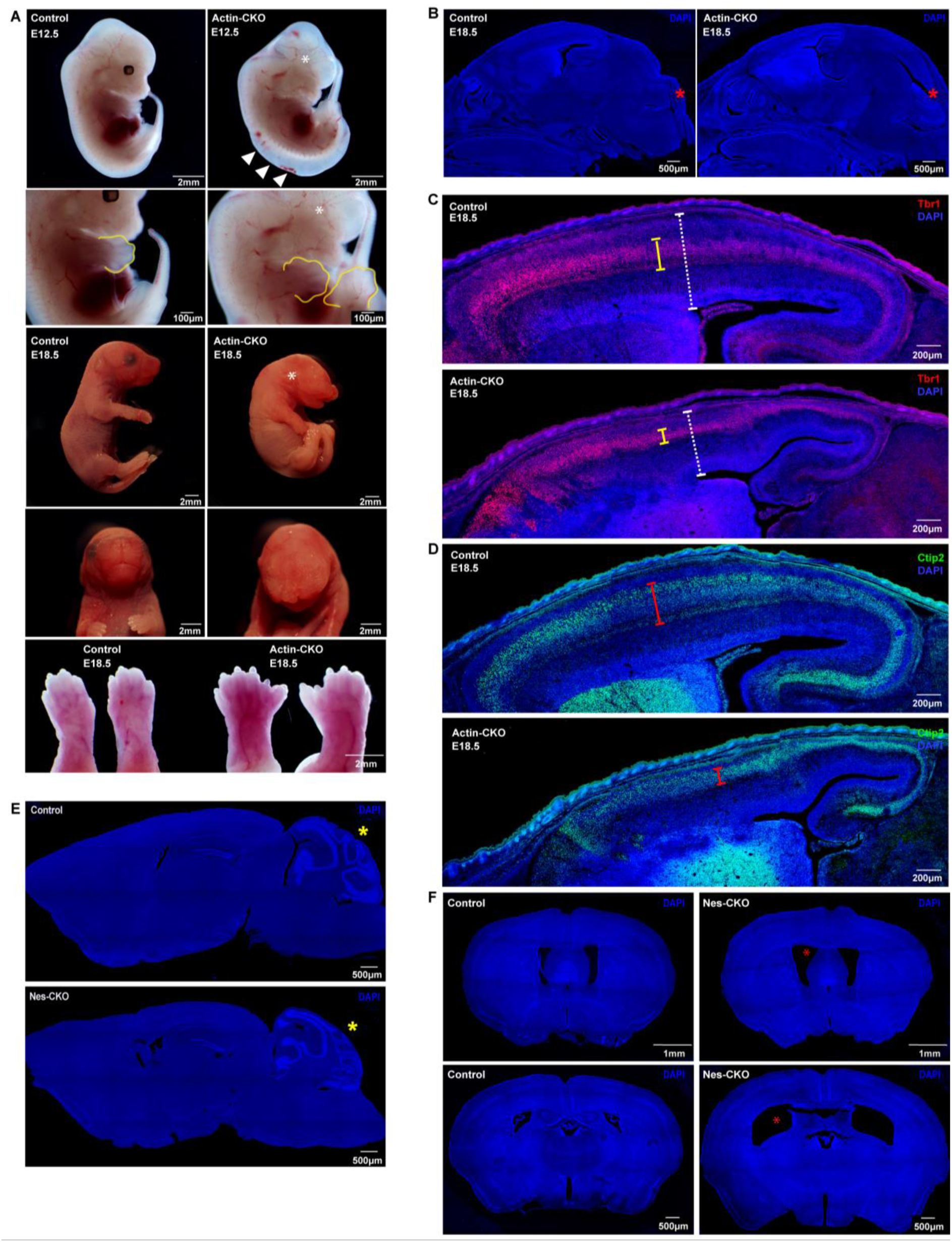
*Rab23*-KO mutants recapitulate the cardinal features of the CS and ciliopathy. **(A)** Representative images show gross morphological appearance of control and actin-CKO mutant mice at E12.5 (top) and E18.5 (bottom), respectively. White asterisks label missing or deformed eyes in actin-CKO mice, white arrowheads show the mis-patterned posterior neural tube. Bottom panel: representative close-up images of the limbs reveal polysyndactyly in actin-CKO mice at E18.5, a similar abnormality is also observed at E12.5 (yellow outline, second panel from top) **(B)** Representative DAPI-stained images depict mid-sagittal sections of the head regions of E18.5 control and actin-CKO mouse embryos. Mis-patterning of the cerebellar anlage (red asterisk) is observed in the actin-CKO embryo. **(C, D)** Representative mid-sagittal brain sections depict mis-patterned and thinned cerebral cortex (white dashed capped arrow) in the actin-CKO mice, which was associated with a thinning of (C) Tbr1+ (yellow capped arrow) and (D) Ctip2+ (red capped arrow) post-mitotic neuron layers. **(E)** Representative DAPI-stained images depict mid-sagittal brain sections of adult control (top) and Nes-CKO (bottom) mice respectively. Cerebellum mis-patterning (yellow asterisk) was observed in adult Nes-CKO mice. **(F)** Representative DAPI-stained images depict coronal brain sections of adult control and Nes-CKO mice, revealing an enlargement of brain ventricles (red asterisks) in Nes-CKO mice. The top panel depicts lateral ventricles at the rostral cortex. Bottom panel depicts lateral ventricles at the rostral hippocampus level.

Furthermore, to investigate the specific neuronal functions of *Rab23*, spatial deletion of *Rab23* was achieved in neural progenitor cells at approximately E10.5 using the Nestin-cre driver line (herein, referred to as Nes-CKO for the homozygous deleted mutant). Nes-CKO mutant mice survived into adulthood, exhibited obesity (data not shown), and showed a significant enlargement of the brain ventricle (Fig. 1F, indicated by red asterisks). However, the morphological appearance of the cerebral cortex in the adult Nes-CKO mutants appeared relatively normal compared to the control counterpart (*Rab23^f/f^*) (Fig. 1E-F).

Consistent with the observed mis-patterned cerebellar anlage in the embryonic actin-CKO mutants, abnormal cerebellar folia formation was also observed in the adult Nes-CKO mice (Fig. 1E, indicated by yellow asterisks). These findings suggest that the mutant mice share several clinical characteristics reported in both Carpenter syndrome and ciliopathies, such as hydrocephaly and obesity (Alessandri et al., 2010; Guemez-Gamboa et al., 2014; Haye et al., 2014; Jenkins et al., 2011; Reiter & Leroux, 2017; Wallmeier et al., 2019; Youn & Han, 2018).

Taken together, the *Rab23* mutants prominently display multiple developmental features that closely resemble both Carpenter syndrome patients and individuals affected by other forms of ciliopathy. This strongly suggests that the loss of *Rab23* function leads to ciliopathy in the mouse mutants. Importantly, the *Rab23* loss-of-function mouse mutants serve as a robust model for Carpenter syndrome, as they successfully recapitulate numerous clinical characteristics observed in human patients. These mutant mice provide valuable animal platforms that will facilitate comprehensive investigations into the underlying disease mechanisms of Carpenter syndrome.

### Rab23 exerts a cell-type specific function in maintaining proper primary cilium formation

Given that the aforementioned features can be found in various ciliopathy disorders (Badano et al., 2006; Damerla et al., 2015; Gerth-Kahlert & Koller, 2018; Liu et al., 2014), as well as Carpenter syndrome (Alessandri et al., 2010; Ben-Salem et al., 2013; Jenkins et al., 2011; Movva et al., 2014), our mouse model data implicates a potential mechanistic association between Carpenter syndrome and ciliopathies. Considering the apparent phenotypic correlation with ciliopathies, we investigate whether the deletion of *Rab23* would disrupt primary cilium formation and/or affect its structural integrity. To visualize primary cilia in various cell types, including epithelial cells, chondrocytes, neuronal progenitors, granule cell precursors (GCP), and mature neurons, we conducted immunohistochemistry staining using specific antibodies against Arl13b and AC3. Consistent with our hypothesis, the mutant mice exhibited a notable decrease in the number of cells in the cerebral cortex bearing primary cilium. This trend was consistently observed in both knockout mutants, i.e. including the Tbr1-expressing cortical intermediate progenitors at embryonic stage (E18.5 of actin-CKO), as well as in NeuN-positive cortical neurons in the adult neocortex of the Nes-CKO mutant (Fig. 2A-B, J-K). Co-immunostaining of adult sagittal brain sections with two ciliary markers i.e. Arl13b and AC3, along with the neuronal marker NeuN, revealed that the majority of cortical neurons in the Nes-CKO mutant neocortex had lost their primary cilia, whereas cortical neurons in the control animals exhibited a higher level of ciliation (Fig. 2J-K). In chondrocytes, although there was no discernible change in the prevalence of ciliation, both the volume and length of primary cilia were significantly reduced (by 17.96 % and 12.98 %, respectively) in the E18.5 actin-CKO mutant, compared to controls (Fig. 2G-I).

**Figure 2.**
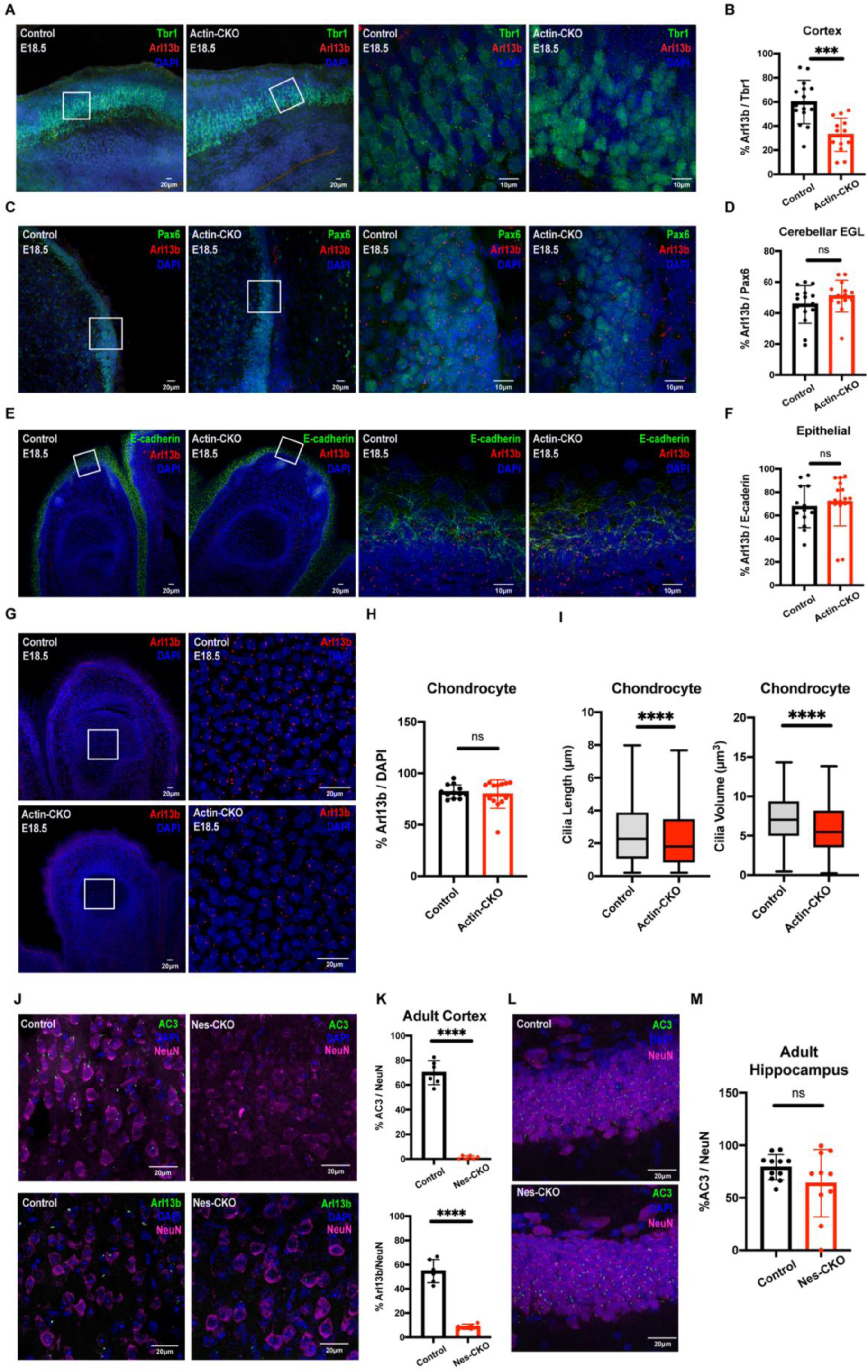
*Rab23* deletion perturbs ciliation in a context-dependent manner. **(A-B)** Representative immunohistochemistry images (low power on left, magnification of boxed region on right) and **(B)** graph depicting quantification of the proportion of Arl13b+ primary cilia against Tbr1+ (green) neocortical layer VI neurons in the neocortex at E18.5. A significant two-fold reduction in the number of primary cilia is observed in the cerebral cortex of actin-CKO mouse embryos. The data represent quantifications from 3 to 4 mice in each genotype. For each animal, 3-4 brain sections of a comparable region were analysed. Each dot represents the percentage count derived from each brain section. Total counts: Control n = 15; actin-CKO n=13 **(C-D)** Representative immunohistochemistry images and **(D)** graph depicting quantification of the proportion of Arl13b+ primary cilia against Pax6+ (green) granule cell precursors in the external granule layer (EGL) of cerebellum anlage at E18.5. The data represent quantifications from 3 to 4 mice in each genotype. For each animal, 3-4 brain sections of a comparable region were analysed. Each dot represents the percentage count derived from each brain section. Total counts: Control n = 15; actin-CKO n=14 **(E-F)** Representative immunohistochemistry images and **(F)** graph depicting quantification of the proportion of Arl13b+ primary cilia against E-cadherin+ (green) epithelial cells lining the epidermal layer at E18.5. The data represent quantifications from 3 to 4 mice in each genotype. For each animal, 3-4 brain sections of a comparable region were analysed. Each dot represents the percentage count derived from each brain section. Total counts: Control n = 13; actin-CKO n=16 **(G-H)** Representative immunohistochemistry images of E18.5 digits and **(H)** graph depicting quantification of the proportion of Arl13b+ primary cilia against chondrocytes residing in the phalanges. The data represent quantifications from 3 to 4 mice in each genotype. For each animal, 3-4 brain sections of a comparable region were analysed. Each dot represents the percentage count derived from each brain section. Total counts: Control n = 12; actin-CKO n=13 **(I)** Graphs depict the measurements of cilia length (left) and volume (right) in the chondrocytes of control and actin-CKO at E18.5, respectively. Boxplots illustrate data from 3 to 4 biological replicates in each genotype. For each sample, the data was collected from 3-4 images of comparable regions of interest. Total number of cilia measured: n = 800-1000 cilia from 3 - 4 biological replicates in each genotype. **(J-K)** Representative immunohistochemistry images and **(K)** graphs depicting quantification of the proportion of AC3+ (top panel green) or Arl13b+ (bottom panel green) primary cilia against NeuN+ (pseudo-colored magenta) neurons in cerebral cortex of control and Nes-CKO mice aged 2-3 months. Similar to actin-CKO mice, neural progenitor cell-specific *Rab23* knockout mutants show dramatically decreased number of ciliated neurons in the cerebral cortex of the animals. **(L-M)** Representative immunohistochemistry images and **(M)** graph depicting quantification of the proportion of AC3+ primary cilia against NeuN+ neurons in the hippocampal CA1 cells of both control and Nes-CKO mice. N = 4 mice in each genotype. For each sample, the percentage of ciliated cells was determined from 3-5 images of comparable regions of interest. ns, not significant, *** *P* value ≤ 0.001, **** *P* value ≤ 0.0001 Unpaired Student’s t-test.

Interestingly, unlike that observed in the cortical layer, the neuronal population in the adult hippocampus CA1 region showed a similar number of AC3-positive primary cilia in both the control and the Nes-CKO mutant group (Fig. 2L-M). Similarly, in the E18.5 actin-CKO mutant, a relatively normal prevalence of ciliation was observed in the Pax6-expressing granule cell precursors (GCPs) in the *Rab23*-deficient cerebellar anlage, as well as in E-cadherin expressing epithelial cells lining the dermal layer (Fig. 2C-F).

In conclusion, these findings represent the first *in vivo* evidence demonstrating that the knockout of *Rab23* specifically impairs the formation and/or maintenance of primary cilia in chondrocytes and specific neuronal populations. This highlights the critical role of Rab23 in primary cilium-dependent functions in the central nervous system (CNS) and bone development in mammals. Interestingly, it is noteworthy that not all neuronal cell types show an evident primary cilia defect upon *Rab23* loss, suggesting that the ciliary function of *Rab23* may be context-dependent and influenced by specific neuronal functions.

### Morpholino knockdown of *rab23* in zebrafish results in selective deciliation of a restricted brain region

Given that the mouse *Rab23* shares 83.13 % sequence identity with zebrafish *rab23* (Supplementary figure 1), we investigated further to find out if the functional effects of *Rab23* on the primary cilium are conserved in the zebrafish model. In order to knock down the *rab23* homolog in zebrafish, we designed a morpholino (MO) oligonucleotide targeting the splicing site at the exon 2-intron 2 border of *rab23* to block proper splicing of *rab23* mRNA. This splicing-blocking MO was expected to cause either complete or partial deletion of exon 2, leading to the production of defective mRNA transcripts. The efficiency and specificity of the splicing blocker MO were assessed using reverse transcription polymerase chain reaction (RT-PCR). Indeed, at 24 hpf following injection of MO, we observed that while the level of internal loading control (i.e. *actin*) remains relatively unchanged (Figure 3B, lower panel, 206 bp), the amount of spliced *rab23* product was greatly reduced, with very little aberrant spliced product detectable (Figure 3B, upper panel). This result suggests that the MO effectively blocked the proper splicing of *rab23*, resulting in a depletion and/or generation of defective *rab23* transcript.

**Figure 3:**
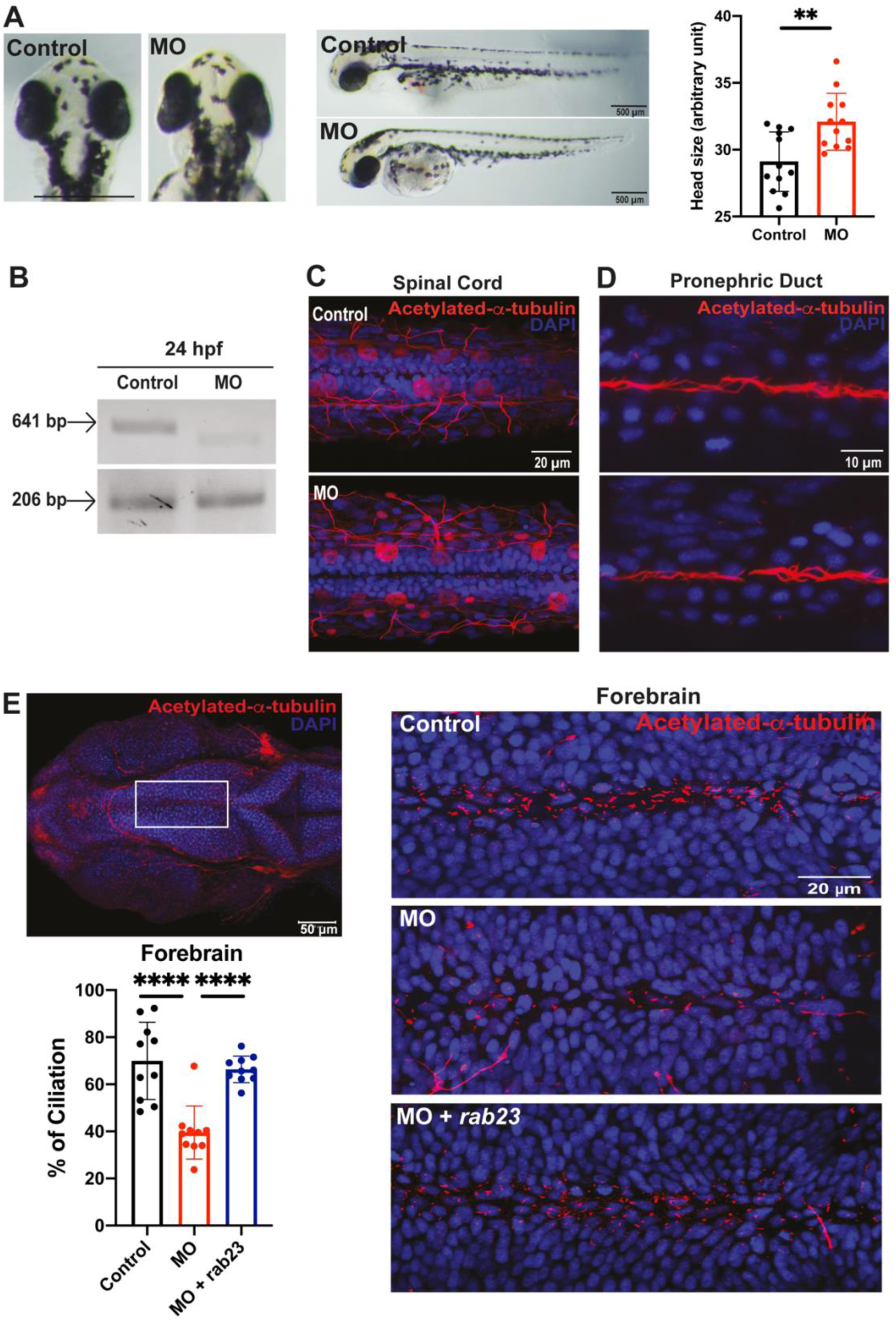
Knockdown of *rab23* in zebrafish affects primary cilia formation at the rostral brain ventricles. **(A)** Morpholino-mediated knockdown of *rab23* in zebrafish. Bright-field dorsal view microscopy images of control and morphant (MO) at 72 hpf. n = 12 for each group, scale bar = 500 *μ* m. Graph representing quantitative head size measurement between control and morphants. The head size was determined by measuring the distance between the eyes in the dorsal view images. The average head size of morphants was slightly yet significantly larger than the control group. ** *P* value ≤ 0.01 Unpaired Student’s t-test. Error bars depict S.D. **(B)** Representative gel image showing normal splicing of *rab23* in control (641 bp) or inhibited splicing in morphants 24 hour-post-fertilization (hpf), resulting in less or shorter spliced product. Actin (206 bp) was used as the internal control. **(C)** Representative images showing largely unaffected cilia number at the central canal of the spinal cord between control and morphant. **(D)** Representative images showing largely unaffected cilia number at the central canal of the pronephric duct between control and morphant. **(E)** Representative images showing an obvious reduction of cilia number present in the forebrain ventricle of morphant as compared to control. Bottom right: rab23 morphant rescued by injecting mRNA of *rab23* (bottom panel). Bottom left panel: Graph presenting quantitative analysis of cilia number in the brain ventricle of control, morphants and rescue group at 24 hpf, respectively. n = 10 for each group. Error bars depict S.D. **** *P* value ≤ 0.0001 One-way ANOVA.

A slight but significant head size difference was observed between the controls and *rab23* morphants, in which the head size of *rab23* morphants was larger than that of controls (Figure 3A). In zebrafish, motile cilia can be commonly found in the pronephric duct and Kupffer’s vesicle, whereas primary cilia can be found along the brain ventricle from the anterior neural tube at the forebrain to the posterior neural tube at the spinal cord. We examined the motile and primary cilia by performing whole-mount immunostaining of acetylated-*α*-tubulin at 24 hpf. Our results showed that there was no significant reduction of motile cilia number in the either pronephric duct (Figure 3D), or the number of primary cilia lining the central canal of the spinal cord (Figure 3C). Interestingly, a prominent reduction in the number of primary cilia was observed in the forebrain ventricle of *rab23* morphants. Quantification revealed that the percentage of ciliated cells in the forebrain ventricle of *rab23* morphants reduced to approximately 40 %, whereas there were about 70 % ciliated cells in the control counterpart (Figure 3E). To rule out potential off-target effects of the morpholino, we performed a rescue experiment by co-injecting MO with *rab23* mRNA and quantified the cilia number (Figure 3E). There was a significant recovery in the number of primary cilia in the rescue group as compared to the morphant without *rab23* mRNA (Figure 3E), confirming that the decrease in the percentage of ciliation was due to the loss of *rab23*.

Thus, in line with the observations in our mouse mutants, silencing *rab23* in zebrafish also resulted in a perturbation of primary cilia in specific neuronal lineages/populations. In particular, our data revealed that zebrafish *rab23* exerts paramount ciliary functional effects in the neurons residing in the forebrain but not in the spinal cord neural tube.

### Neural progenitor cells and neurons differentiated from Carpenter syndrome patient iPSCs exhibit impaired formation of the primary cilium

We then investigated whether *RAB23* also plays a role in regulating the primary cilia formation in neuronal cells of human origin. For this purpose, we obtained patient-derived skin fibroblast cells carrying a biallelic p.(L145*) nonsense mutation in the *RAB23* gene (Jenkins et al., 2007, 2011). These fibroblast cells were reprogrammed into induced pluripotent stem cells (iPSC) clones. The successfully reprogrammed iPSC clones exhibited a normal karyotype profile, and their pluripotency was confirmed by abundant expression of human stem cell markers SSEA4, NANOG, OCT3/4, and TRA-1-60 (Figure 4A-C, images represent results of patients-iPSCs with p.(L145*) mutation).

**Figure 4.**
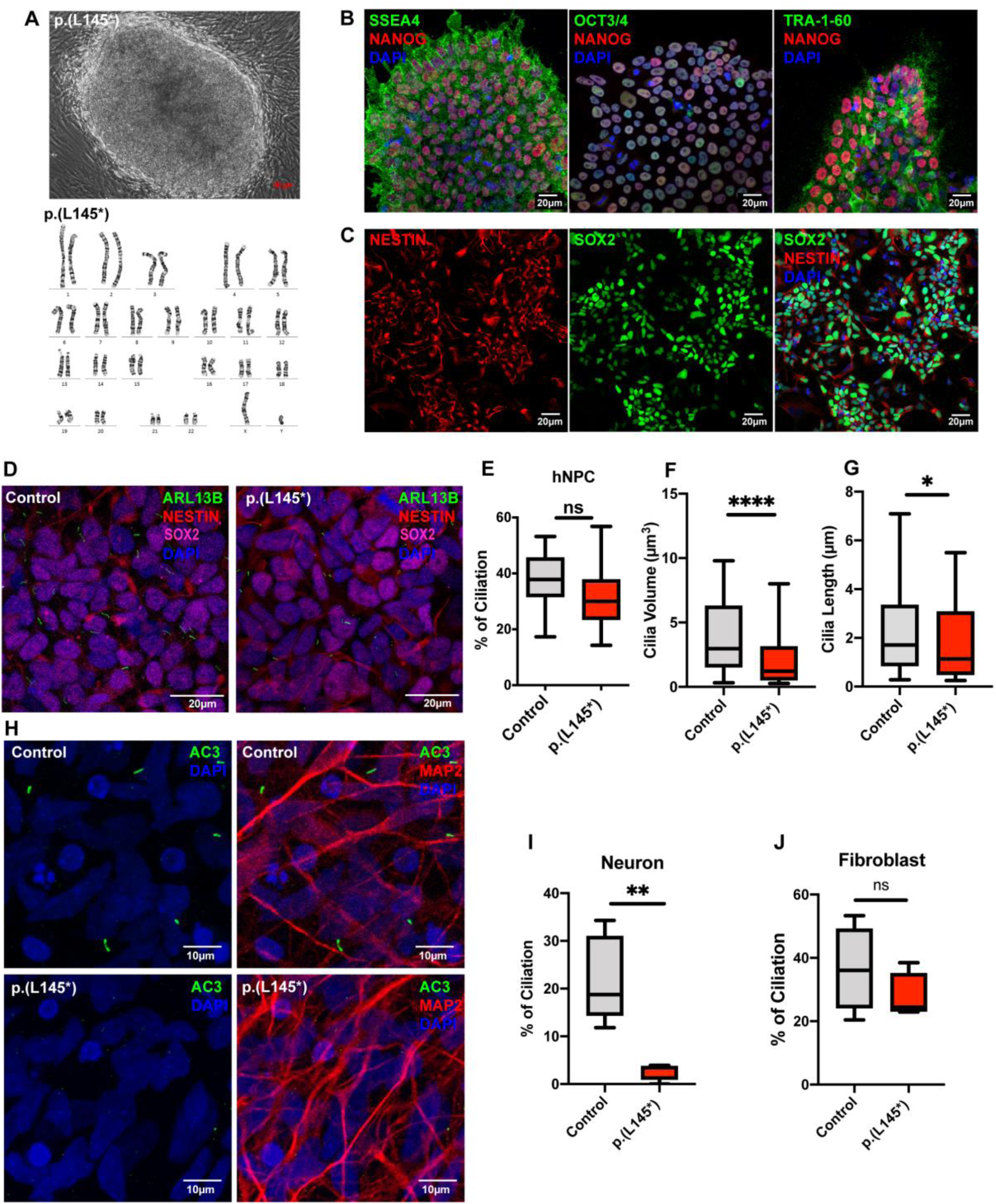
CS patient-derived iPSC-differentiated neurons exhibit impaired ciliogenesis. **(A-B)** Fibroblasts isolated from skin biopsy from a CS patient bearing the *RAB23* biallelic p.(L145*) nonsense mutation was reprogrammed into iPSC clones. **(A)** Top : A representative phase contrast image showing successfully reprogrammed-iPSC clone displaying a normal karyotype (bottom), and pluripotency characterized by positive **(B)** co-immunostainings of human stem cell markers SSEA4, NANOG, OCT3/4 and TRA-1-60 respectively. **(C)** Representative co-immunocytochemistry images depicting positive expression of NESTIN (red) and SOX2 (green) on neural progenitor stem cells induced from iPSCs. **(D)** Representative co-immunocytochemistry images depicting healthy adult and CS patient iPSC-derived human neural progenitor cells (hNPCs) co-immunostained for SOX2 (magenta), NESTIN (red) and primary cilia marker ARL13B (green). **(E)** Graph depicts the quantification of the percentage of ciliation in hNPCs. Box plot represents data from four independent experiments. n.s.= not significant. Unpaired Student’s t-test. **(F-G)** Box plots depict the measurements of (F) cilia length and (G) cilia volume on hNPCs. Data represents the measurements of ∼65–80 cilia in each genotype obtained from three independent experiments. **** *P* value ≤ 0.0001, \**P* value ≤ 0.05 Unpaired Student’s t-test. **(H-I)** Representative co-immunocytochemistry images depict MAP2+ (red) neurons differentiated from iPSCs of healthy adults (control) and CS patients (p.(L145*)). Primary cilia were labelled by AC3 (green). **(I)** Graph depicts the percentage of ciliation, i.e. quantification of the proportion of ciliated neurons (AC3+ MAP2+) in the MAP2 positive neuronal population. A significant reduction in ciliation was observed in neurons bearing p.(L145*) mutation. Box plot represents data from five independent experiments. ** *P* value ≤ 0.01 Unpaired Student’s t-test. **(J)** Graph depicts the quantification of the percentage of ciliation in primary fibroblast cells. The primary cilia in human fibroblast were visualised by immunostaining of acetylated-alpha-tubulin and ARL13B. Primary fibroblasts cultured from a healthy donor and a CS patient with p.(L145*) mutation show comparable percentages of ciliation. Box plot represents data from four independent experiments. n.s.= not significant. Unpaired Student’s t-test.

We examined the primary cilia in patient iPSC-induced neural progenitor cells (hNPC), iPSC-differentiated neurons, and patient fibroblasts. Normal fibroblasts and iPSCs reprogrammed from healthy human dermal fibroblasts were used as the respective controls. Interestingly, the patients’ neural progenitor cells exhibited a significant reduction in the volume and length of the primary cilium axoneme, despite an unaffected percentage of ciliation (Figure 4D-G). On the other hand, the mutant neurons exhibited a significantly reduced ciliation prevalence (reduced by 88.26 %, Figure 4H-I). No discernible cilia defects were observed in the patient fibroblasts (Figure 4J). These findings indicate that *RAB23* plays an important role in human neuronal ciliogenesis, and the loss of *RAB23* function causes a ciliopathy in humans.

### Loss of *Rab23* impairs primary cilium-dependent Sonic Hedgehog signaling pathway activation in the neural progenitor cells

We hypothesize that the decrease in ciliation disrupts the signal transduction of primary cilium-dependent Shh signaling pathway, potentially playing a role in the phenotypical abnormalities and developmental defects observed in the *Rab23*-KO mutant and Carpenter syndrome patients. To investigate this, we examine the ciliation prevalence, and measure the response of *Rab23*-KO cortical neural progenitor cells (NPCs) to the stimulation of the Shh signaling pathway in primary culture. Consistent with the *in vivo* findings in the cerebral cortex of the E18.5 actin-CKO mutant (Figure 2A), an apparent reduction in the percentage of ciliated cells was observed in the *Rab23*-KO primary cortical NPCs culture isolated from Nes-CKO mutant (Figure 5A-B). Additionally, akin to the observations in the human NPCs derived from the CS patient iPSC, the cortical NPCs cultured from *Rab23*-KO mice also exhibit a shortened cilia length and reduced cilia volume (Figure 5B).

**Figure 5.**
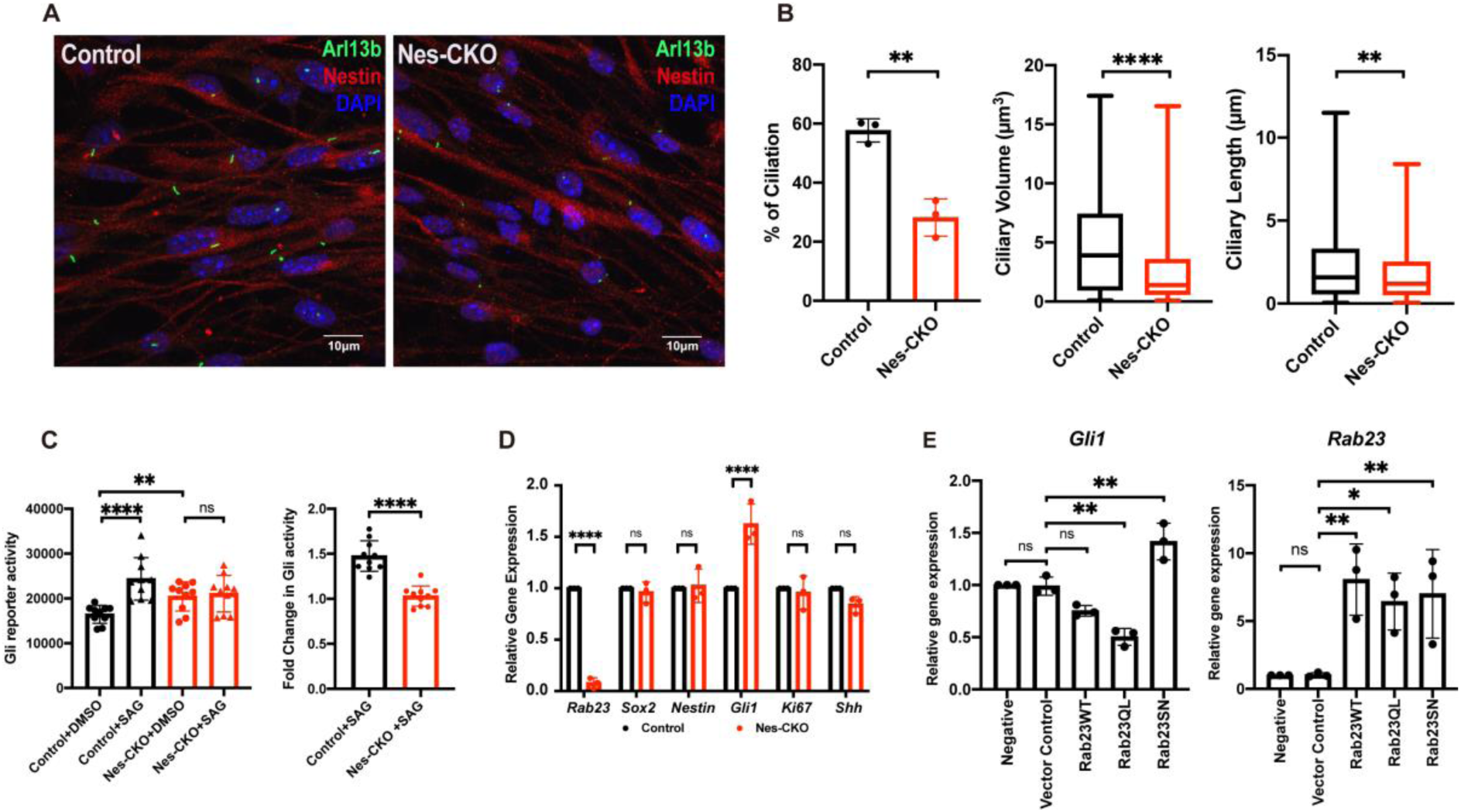
*Rab23*-deleted murine neural progenitor cells show disrupted ciliation and compromised response to Smo-dependent Hedgehog signaling pathway activation. **(A)** Representative immunocytochemistry images depict control and Nes-CKO cortical NPCs immunostained for Nestin (red) and primary cilia marker Arl13b (green). **(B)** Graphs depict the quantification of the percentage of ciliation, ciliary volume and length in mouse cortical neural progenitor cells culture. Each dot represents the average percentage value of each independent experiment. * *P-*value ≤ 0.05, ** *P-*value ≤ 0.01, **** *P*-value ≤ 0.0001, Unpaired Student’s t-test. For cilia volume and length measurement; Control n = 230, Nes-CKO n = 211, measured from three independent experiments. **(C)** Graphs depict the (left) 7Gli:GFP reporter signals and the (right) relative fold-change in control and Nes-CKO cortical NPCs treated with 200 nM SAG and DMSO respectively for 24 hours. Data points represent triplicate readings from three independent experiments. Left: **** *P*-value ≤ 0.0001, ** *P*-value ≤0.01, n.s.= not significant, two-way ANOVA. Right: **** *P*-value ≤ 0.0001, Unpaired Student’s t-test. **(D)** Graph depicts the relative gene expression level of control and Nes-CKO cortical NPCs quantified by real-time qPCR. **** *P*-value≤0.0001, n.s.= not significant, two-way ANOVA. **(E)** Graphs depict the relative gene expression levels of *Gli1* and *Rab23* in wild-type cortical NPCs overexpressing wild-type and mutant forms of Rab23 at day 6 post-transduction. NPCs were transduced with lentiviruses carrying different overexpression construct i.e. Rab23WT, Rab23Q68L, Rab23SN respectively. The gene expression levels of *Gli1* and *Rab23* at day 6 post-transduction were quantified by real-time qPCR. ** *P-*value ≤ 0.01, n.s.= not significant, one-way ANOVA.

After confirming the structurally defective primary cilia in *Rab23*-deleted NPCs culture, we employed SAG, a Smoothened (Smo) agonist, to evaluate the sensitivity of the NPCs to the primary cilium-dependent activation of Shh signaling pathway. SAG triggers the Shh signalling pathway by promoting the Smo translocation to the primary cilium axoneme, thereby activating the primary cilium-dependent Shh signaling pathway. Therefore, a dysfunctional primary cilium would render the cell becoming unresponsive to SAG stimulation. A green fluorescence protein (GFP) Gli reporter system was used to determine the level of Shh signalling pathway activation based on the level of Gli activity.

In line with our speculation, *Rab23*-KO neural progenitor cells did not exhibit responsiveness to the SAG stimulation, showing no discernible change in the Gli reporter activity level following a 24 hour stimulation with 200 nM SAG. Conversely, the control (*f/f*) mouse neural progenitor cells demonstrated a significantly increased level of Gli reporter activity as compared to the DMSO vehicle control group, indicating a robust response and activation of the Shh signaling pathway (Figure 5C). These findings suggest that the loss of *Rab23* impairs the NPCs’ response to primary cilium-dependent Shh signaling pathway activation due to dysfunctional primary cilia.

Notably, in the absence of SAG stimulation, *Rab23-*KO NPCs at basal level (DMSO) showed an up-regulated Gli activity compared to the control group (Figure 5C, Control+DMSO versus Nes-CKO+DMSO). This observation is consistent with our previous observations in cerebellar granule progenitor cells (C. H. Hor & Goh, 2019; C. H. H. Hor et al., 2021), and aligned with the well-established negative regulatory role of *Rab23* in the Shh signalling pathway (Eggenschwiler et al., 2001). Consistent with the Gli reporter assay results, qPCR analysis of gene expression also revealed ectopic activation of the Shh signaling pathway in the mutant NPCs, as evidenced by elevated *Gli1* expression (a downstream transcriptional readout of Shh signaling activity) (Figure 5D). Interestingly, despite the elevated *Gli1* expression, the expression levels of *Shh,* NPC markers i.e. *Nestin* and *Sox2,* and the cell proliferation marker *Ki67,* remained unchanged (Figure 5D). These results may suggest a partial activation of the Shh signaling pathway following the deletion of *Rab23* in NPCs.

While Rab23 is known for its role in modulating the Shh signaling pathway, it has remained unclear whether this function requires its GTP/GDP cycling-dependent property, which is a classical molecular characteristic of small GTPases. To address this, we overexpressed wild-type full-length Rab23, constitutively active (GTP-bound) form of Rab23, i.e Rab2368QL, and dominant negative (GDP-bound) form of Rab23, i.e. Rab23S23N in wild-type NPCs, respectively. The *Gli1* expression levels were then measured and compared between these overexpression groups. The data showed that the overexpression of Rab23Q68L in wild-type mouse cortical NPCs led to a decrease in the expression of *Gli1*. Conversely, overexpressing the dominant negative (GDP-bound) form of Rab23, specifically Rab23S23N, resulted in an increase in *Gli1* expression (Figure 5E), similar to the observations in *Rab23*-Nes-CKO NPCs (Figure 5D). These results indicate that Rab23 modulates Shh signaling activity through its classical GTP/GDP cycling-dependent functions, presumably involving its specific guanine nucleotide exchange factor (GEF) and GTPase-activating protein (GAP).

## Discussion

Our study underscores the critical and evolutionarily conserved functions of *RAB23* in preserving morphologically intact and functional primary cilia across various vertebrates, including the zebrafish, mice, and humans, notably in a cell type distinctive manner (Figure 6). This study unveils that conditional knockout mouse mutants of *Rab23* exhibit a spectrum of phenotypic and pathological traits reminiscent of the classical clinical features reported in both Carpenter syndrome and ciliopathy disorders in humans. Consistent with the phenotypic association to ciliopathy, examination of primary cilia in the *in vivo* tissue section samples of E18.5 and adult *Rab23* conditional KO mice revealed profound ciliary anomalies in several cell types such as the neocortical postmitotic and mature neurons, neural progenitor cells, and chondrocytes. In resonance with the structurally perturbed primary cilia, *Rab23-*KO mouse neural progenitor cells also displayed dysfunctional primary cilia, as evidenced by the desensitised response to the primary cilium-dependent activation of the Hh signaling pathway (Fig. 5C). These results further suggest that dysfunctional primary cilia, to some extent, may underlie the pathological presentation of Carpenter syndrome.

**Figure 6.**
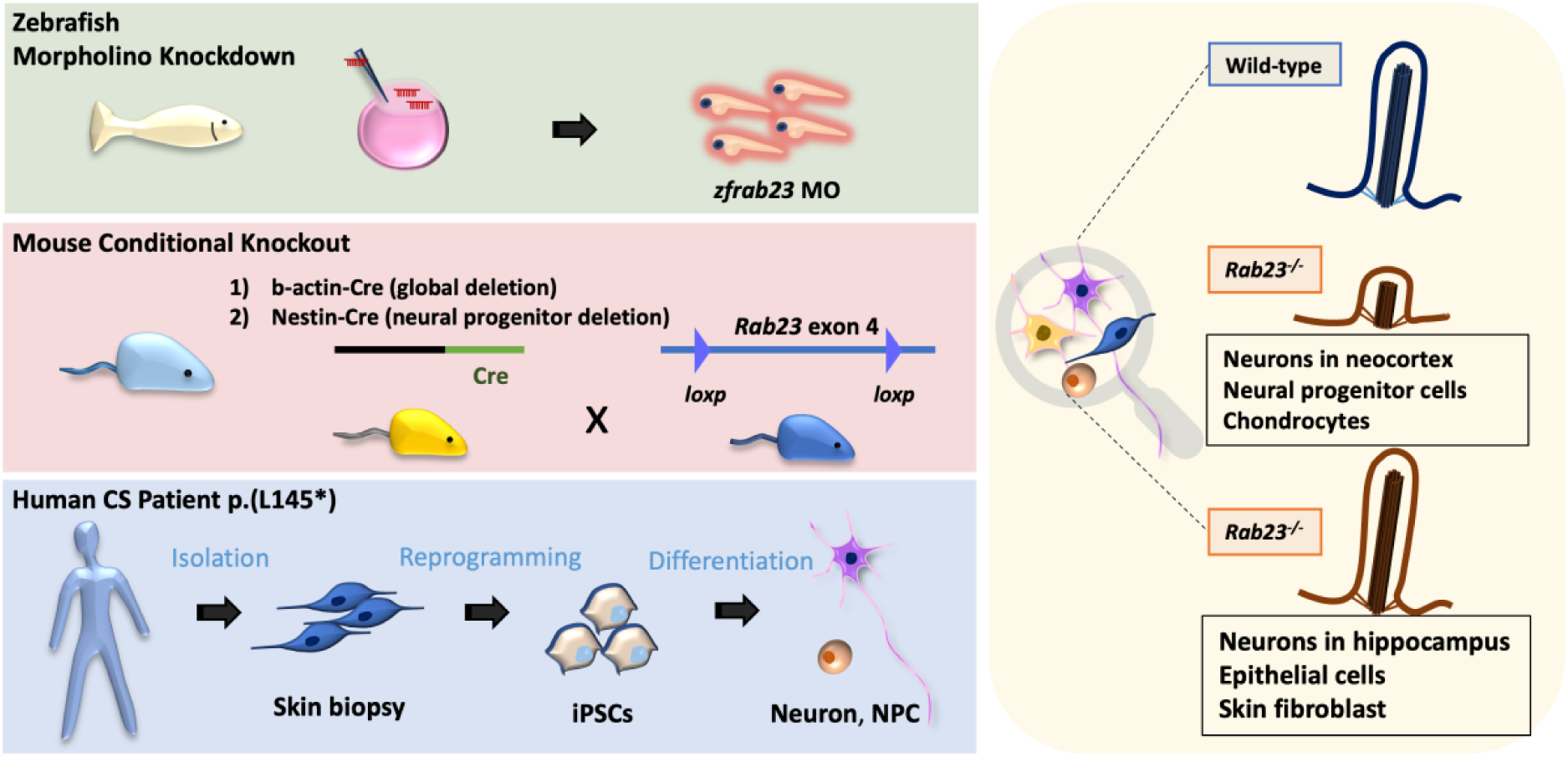
Schematic illustration of the conserved functions of Rab23 in primary cilia. *Rab23* exerts evolutionarily conserved and context-specific functions in maintaining morphologically intact and functional primary cilia. Specifically, in the absence of *Rab23*, perturbations in ciliogenesis were found in distinct neuronal populations in the neocortex, neural progenitor cells and chondrocytes.

Considering the important roles of Rab23 in ciliary trafficking, earlier studies have speculated on the potential connection between Rab23 and ciliopathy (Boehlke et al., 2010; C. H. Hor & Goh, 2019; Leaf & Von Zastrow, 2015; Lim & Tang, 2015; Zhao et al., 2023). However, as far as our current knowledge extends, there remains a lack of direct evidence substantiating the causal link between *RAB23* mutation and Carpenter syndrome with ciliopathy. Our data reveals for the first time that Carpenter syndrome patients (with biallelic *RAB23* p.(L145*) mutation)-derived iPSCs differentiated neurons and neural progenitor cells exhibit primary cilia anomalies. Moreover, we also show that different vertebrate models of *RAB23* loss-of-function mutants consistently presented with several pathological phenotypes and cellular characteristics typical of human ciliopathies. Our work thus provides compelling and direct evidence demonstrating that *RAB23* mutation is clinically linked to ciliopathy, further validating the classification of Carpenter syndrome as one of the disorders within the ciliopathy spectrum.

Intriguingly, in the absence of *Rab23*, the perturbations of primary cilium appeared to selectively affect specific cell types and neuronal populations. This cell-type-specific phenomenon was consistently observed in both *in vivo* mouse and zebrafish models, as well as the CS patient-derived cells. Importantly, our findings on the context-dependent ciliary function of *Rab23* help to explain the discrepancies previously noted in independent studies where primary cilium defects were reported in some experimental models but not in others. (Fuller et al., 2014; Gerondopoulos et al., 2019; Leaf & Von Zastrow, 2015). To examine the structural damage and/or the loss of primary cilia, various primary cilium markers were employed for cilia examinations in this study. For example, ciliary membrane markers including ARL13B and AC3, along with the ciliary microtubule marker, acetylated-α-tubulin, were utilized to visualize the primary cilia axoneme in different cell and tissue samples.

Our results appear to differ from prior findings on *Rab23^-/-^*null mutant mice, where embryos at the 0-1, 3-4, and 5-6 somite stages displayed morphologically normal nodal cilia, despite exhibiting left-right patterning defects (Fuller et al., 2014). Notably, previous studies did not investigate the ciliary functions of *Rab23* at later developmental stages because the *Rab23^-/-^* null mutant embryos in the C3Heb/FeJ background did not survive beyond E13.5 (Eggenschwiler et al., 2001; Fuller et al., 2014). Here, we employed a Cre-loxp transgenic knockout approach, i.e. using actin-Cre to drive the global deletion; and Nestin-Cre for neural progenitor cells-specific deletion of *Rab23* in the C57/BL6 background. These approaches have extended the viability of the mutants up to postnatal day 0 and the adult stage, respectively, allowing us to delineate the important ciliary functions of *Rab23* during late developmental and adult stages.

In these mutant mice, our results indicate that *Rab23* plays a pivotal ciliary function in the neocortical region but not in the hippocampal neurons. A notable decrease in ciliation prevalence was observed in both the Tbr1-expressing postmitotic neurons of the neocortex at late embryonic stages and in the NeuN-expressing mature neocortical projection neurons at adult stages. These findings unveil a previously underappreciated function of *Rab23* in the adult central nervous system. Consistent with this finding, neurological-associated disorders such as intellectual disability and schizophrenia have been reported in cases of CS (Alessandri et al., 2010; Bersani et al., 2003; Lodhia et al., 2021). Our earlier work also revealed the requirement of *Rab23* in modulating the migration of neocortical projection neurons during embryonic brain development (C. H. H. Hor & Goh, 2018). Given these apparent correlations, further investigations to understand the molecular and cellular roles of Rab23 in neuronal development and neuronal pathophysiology are warranted.

In a similar vein, a previous study in the zebrafish model showed that morpholino-induced *rab23* knockdown morphants exhibited morphologically normal cilia in Kupffer’s vesicle at the 8-somite stage, albeit with disrupted left-right patterning (Fuller et al., 2014). In our current investigation, we delved deeper to examine primary cilia in several tissue regions of *rab23* zebrafish morphants at a later embryonic stage, specifically at 24 hours post-fertilization. Our findings highlighted a significant disturbance in the quantity of primary cilia along the ventricular lining of the forebrain neural tube. Conversely, primary cilia in other regions such as the pronephric duct and spinal cord neural tube appeared normal and were indistinguishable from the control counterpart.

We further demonstrate that Rab23 exerts GDP/GTP-binding dependent regulation of the Hedgehog signaling pathway in the neural progenitor cells. Compared to the control and the wild-type Rab23 overexrepression group, the neural progenitor cells overexpressing the GTP-bound constitutively active form of Rab23 exhibit further inactivation of the Hh signaling pathway, whereas overexpression of the GDP-bound constitutively inactive mutant of Rab23 leads to the ectopic activation of the Hh signaling pathway. These results indicate that the classical Rab GTPase properties of Rab23, including its guanine nucleotide exchange factor (GEF) and the GTPase-activating protein (GAP)-dependent on/off actions, are essential for dynamically modulating the Hh signaling pathway in neural progenitor cells.

It’s worthy to note that Carpenter syndrome presents a distinct phenotypic spectrum compared to other ciliopathies such as Joubert syndrome, Bardet-Biedl syndrome and AlstrÖm syndrome. Based on our data, it’s plausible that this unique phenotypic spectrum may be attributed to the context-specific ciliary roles of Rab23, as well as the ciliary-independent regulatory functions of Rab23 on cell migration and other developmental signalling pathway such as the FGF, ERK and nodal signaling pathways (Fuller et al., 2014; Hasan et al., 2020; C. H. H. Hor & Goh, 2018). For instance, it was reported that Rab23 influences FGFR and ERK1/2 signaling pathways in osteogenesis (Hasan et al., 2020).

Our previous study (C. H. H. Hor et al., 2021) and our current findings demonstrate that Rab23 exerts bi-directional functions in regulating Hedgehog signaling activity. It can positively influence the pathway through promoting primary cilia-dependent Hh signal transduction (Figure 5A-C), and negatively affect it by acting on Gli transcription factors in the cytoplasm (Figure 5E). This indicates that different tissues and cell types may respond differently to the loss of Rab23 in terms of Hedgehog signaling activity, whereby Rab23-ablated cells without primary cilia may show partial activation of Hedgehog signaling due to the diminished primary cilia-dependent signaling response, whereas those with intact primary cilia may exhibit ectopic activation of the pathway. These findings suggest that Rab23 plays critical roles in modulating a context-dependent differential regulation of Hedgehog signaling pathway.

Our comprehensive *in vitro* and *in vivo* evidence strongly suggests that *RAB23* plays instrumental roles in ciliary function, thus positioning *RAB23* as part of the family of ciliary genes. Collectively, our findings provide new insights into the pathological mechanisms of Carpenter syndrome and pave the way for future development of clinical interventions. Additionally, the mouse mutants represent a valuable animal model for further investigations into the disease mechanism of CS. Continued research in this area is imperative to advance our understanding and improve the clinical management of debilitating ciliopathy disorders.

## Materials and Methods

### Animal and ethics approval

All animal protocols followed animal handling guideline approved by HKBU Institutional Research Ethics Committee, the Department of Health, Hong Kong, as well as IACUC Singhealth, Singapore. The *Rab23-flox* mouse described in our previous work(C. H. H. Hor et al., 2021), was generated by Ozgene Pty Ltd. Nestin-Cre (The Jackson Laboratory cat. no. 003771) was a kind gift from Shawn Je H.S. from Duke-NUS Medical School. β-actin-Cre driver line was purchased from The Jackson Laboratory. Mice were housed in the animal facility, Department of Chemistry, HKBU, and the Specific Pathogen Free (SPF) animal facility at Duke-NUS Medical School, Singapore. The animal experiments described in this project included unbiased data from both female and male mice unless otherwise specified. The control animals were heterozygous *Rab23^f/+^*or homozygous *Rab23^f/f^* mutants.

### Ethical approval for human clinical samples

Written consent was obtained from the parents of the CS patient before the collection of the skin fibroblast sample for this study. IRB approval was provided by Central Oxford Research Ethics Committee, ref. C02.143, study name ‘*Genetic basis of craniofacial malformations’.* Studies conducted on the same CS patient were previously reported in Jenkins et al 2007 (Jenkins et al., 2007) and Perlyn & Marsh 2008 (Perlyn & Marsh, 2008).

### Cryosection and immunohistochemistry

The cryosection and immunohistochemistry staining protocols have been reported in our previous work (C. H. H. Hor et al., 2021). In brief, immediately post-perfusion, adult mouse brain samples were dissected from the skull and post-fixed in 4% paraformaldehyde for 2 hours on ice. For embryonic mouse samples, tissues of interest were dissected in ice-cold phosphate-buffered saline (PBS) immediately upon decapitation, transferred to 4% paraformaldehyde and incubated for 1 - 4 hours on ice for fixation. Fixed samples were transferred to 30% sucrose in phosphate buffer and stored at 4 °C until subjected to cryosection. Tissue samples were sliced at 20 µm thickness and mounted on Superfrost® glass slides. For immunohistochemistry staining, samples were subject to microwave heated antigen retrieval, 1 hour blocking in blocking buffer (1% bovine serum albumin (BSA), 2% donkey serum and 0.3% Tx-100) at room temperature, and primary antibody incubation at 4 °C overnight. On the next day, samples were subjected to 3 times of PBS washing for 15 mins each, followed by secondary antibody incubation for 2 hours at room temperature. Post-incubation, samples were washed 3 times with PBS for 15 mins in each wash and mounted with home-made mounting media in semi-air-dry conditions. The primary antibodies and dilution factors used were: anti-rabbit Arl13b (Proteintech, 17711-1-AP 1:1500), anti-mouse Arl13b (DSHB/NeuroMAB clone N295B/66 1:800), anti-rabbit AC3 (Santa Cruz discontinued 1:1500), anti-mouse Nestin (Sigma/Millipore MAB5326 1:1000), anti-mouse Map2 (Sigma, M9942 1:1000), anti-rabbit Pax6 (Biolegend/Covance, PRV-278P 1:1000), anti-mouse NeuN (Sigma/Millipore, MAB377 1:800), anti-rabbit Tbr1 (Abcam, AB31940 1:500), anti-mouse acetylated alpha tubulin (Sigma Aldrich, T6793 1:2000). Secondary antibodies were from Invitrogen Life Technologies, Alexa Fluor®, used at 1:1000 diluted in blocking buffer. Confocal images were captured by Nikon Confocal and Carl Zeiss LSM 710 confocal microscope.

### Reprograming of human fibroblast to iPSCs

The human fibroblast reprogramming protocol was adapted from Okita *et. al.* (Okita et al., 2011) and Su *et. al.* (Su et al., 2015). Briefly, human patient dermal fibroblasts were co-electroporated with episomal vectors (pCXLE-hOCT3/4, shp53, pCXLE-hSK, pCXLE-hUL) at 1 µg each. Electroporated cells were immediately seeded on Matrigel (BD Biosciences #354277)-coated 6 well plates and refreshed with mTeSR1 (Stem Cell Technologies) medium daily for approximately one month until the iPSC colonies reached the optimal size. At optimal size, the colonies were isolated, expanded, and characterised for normal karyotype and pluripotency through immunostaining examination of stem cells markers expression, including NANOG, TRA-1-60, OCT3/4 and SSEA4.

### Human NPC induction and neuron differentiation from iPSCs

The human NPC induction protocol was modified from Li *et. al.* (Li et al., 2011) and Su *et. al.* (Su et al., 2015). Briefly, iPSCs at 20% confluency were cultured in NPC induction media (same as that used in Su et al., 2015) for 7 days, which was. replaced with fresh media every other day. On day 7, the culture was ready for passage and expansion into a 20 µg/ml PDL (Sigma Aldrich #P0899) and Matrigel (BD Biosciences #354277)-coated dish. Subsequently, the NPC culture was maintained in NPC maintenance media (same as that described in Su et al., 2015).

For the neuronal differentiation, NPCs were seeded at 5 x 10^3^ cells/cm^2^ in 20 µg/ml PDL (Sigma Aldrich #P6407) and Matrigel coated coverslips in 24 well plate with neuronal differentiation medium (1x N2 Supplement (Life Technologies #17502048), 1x L-Glutamine (Life Technologies #25030081), 1x Pen/Strep (Life Technologies #15140122) in DMEMF12/Neurobasal (Life Technologies #21103049) medium mixture at 1:1 ratio), with. 10 µM of Rho-associated protein kinase (ROCK) inhibitor (Y-27632, ATGG) freshly added. One-half of the media was gently refreshed every 2-3 days. MAP2-expressing differentiated neurons were obtained on day 21-28.

### Mouse neural progenitor cell culture and Smoothened agonist (SAG) stimulation

Postnatal day 0 cerebral cortical tissues were dissected and digested in digestion buffer consisting of Earle’s Balanced Salt Solution (EBSS) containing Papain (Worthington LS003126) diluted at 1000x, 0.1 mg/ml DNaseI (Roche catalog #11284932001), 5.5 mM cysteine-HCl, incubated for 30 min at 37°C. The digested tissues were then resuspended and dissociated into single cells suspension by gentle pipetting with a P1000 pipettor in DMEM/F-12 medium. The cell suspension was passed through a 70-μm cell strainer (SPL catalog #93070) and centrifuged at 1,200 rpm for 5 min. The cell pellets were then resuspended and cultured in DMEM/F12 maintenance medium containing 20 ng/ml FGF, 20 ng/ml EGF, 2 µg/ml Heparin, 1% N2 Supplement (Life Technologies, catalog #17502048) and 1% penicillin/streptomycin. The cells were cultured on a non-adherent surface for 5-7 days to allow neurosphere formation. When the neurospheres reached a diameter of 120-150 µm, the medium with neurospheres was transferred to a conical centrifuge tube and centrifuged at 1,000 rpm for 5 min. The neurospheres were incubated in Accutase for 5 min, and then maintenance medium was added to terminate the enzyme activity and dissociate the neurospheres into a single-cell suspension. Dissociated NPCs were plated on 20 µg/ml poly-D-lysine (Sigma-Aldrich, catalog #P0899)-coated culture plates. Smoothened agonist (SAG) stimulation was performed as previously reported (C. H. H. Hor et al., 2021). Briefly, the NPCs were seeded at equal cell density at 3 × 10^4^ per well on a 96-well plate and infected by lentivirus carrying a 7Gli:GFP reporter construct (Addgene #110494; Li et al., 2016). On day 4 post-infection, 0.2 μM SAG (Cayman Chemical, catalog #11914-1) was added to the culture and incubated for 24 hour. Green fluorescence reporter signals were measured by fluorescence microplate reader 24 hour post-treatment. For the control group, an equal volume of dimethyl sulfoxide (DMSO) was added as the negative control to drug treatment.

### Lentivirus packaging and production

The lentiviruses used for the overexpression of full-length Rab23 and its mutant forms in the mouse NPCs were produced using HEK293GP packaging cells (Takara) co-transfected with VSV-G, delta8.9, together with FUGW-GFP as the lentiviral vector carrying the Rab23WT, Rab23S23N, and Rab23Q68L, respectively. To harvest the viruses, the culture media was collected 24 hour post-transfection, concentrated through standard ultra-centrifugation methods, and store in aliquots at -80 ℃. The FUGW constructs inserted with the Rab23 full-length and the mutant forms used were described in our previous work (C. H. H. Hor et al., 2021). In brief, Rab23S23N is the constitutive negative mutant, whereas Rab23Q68L is the GTP-bound constitutive active mutant.

### Real-time qPCR

Total RNA was extracted using Trizol (Vazyme). cDNA was reverse transcribed from equal amounts of RNA using the iScript™ cDNA Synthesis Kit (Biorad). Using equal volumes of cDNA as the PCR template, the corresponding gene expression levels were evaluated by quantitative real-time PCR (qPCR) using a TB Green Master Mix (Takara). PCR primers used were:

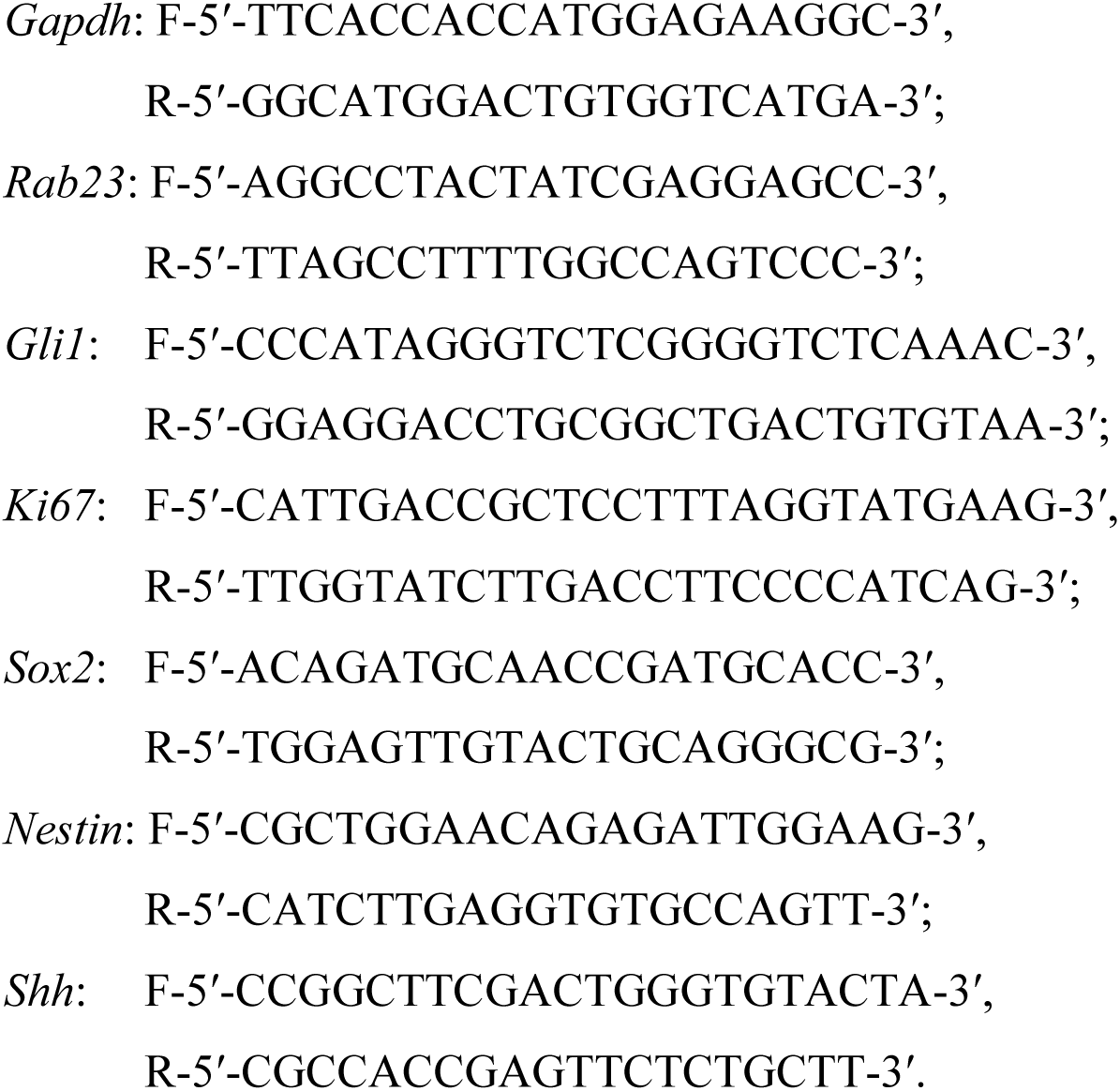

### Immunocytochemistry

Cells cultured on coverslips were fixed with 4% paraformaldehyde for 15 minutes at room temperature. Subsequently, the cells were washed with 1 x PBS and permeabilized with 0.1% Triton X-100 for 15 minutes and blocked in a blocking solution of 1% BSA in PBS for 1 hour at room temperature. The cells were then subjected to overnight incubation at 4 °C with primary antibodies diluted in the blocking solution. Primary antibodies used were: mouse anti-Nestin (1:1000, Santa Cruz Biotechnology), rabbit anti-Arl13b (1:1000, Proteintech), mouse anti-OCT3/4 (1:500 Santa Cruz Biotechnology), rabbit anti-NANOG (1:100, Cell Signaling Technollogy), mouse anti-SSEA4 (1:500, Milliporer), mouse anti-TRA-1-60 (1:100, Santa Cruz Biotechnology), goat anti-SOX2 (1:500 Santa Cruz Biotechnology). After three washes with PBS, the cells were incubated with fluorophore-conjugated secondary antibodies (Alexa Fluor, Life Technologies) for 1 hour at room temperature. Following three additional PBS washes, the cells were mounted in homemade mounting media, kept in dark and air-dry before proceeding to confocal imaging.

### Injection of zebrafish

Wild-type zebrafish of the AB strain were maintained under standard conditions of fish husbandry. Freshly fertilized zebrafish eggs were injected with mRNA (100 ng/μl) and *rab23* morpholino (600 μM) at the one- to two-cell stage in a volume of approximately 1 µl. Approximately 200 embryos were injected for morpholino knockdown or mRNA rescue. The injected embryos were cultured at 28 *^°^*C, and embryos were harvested at respective timepoints for RT-PCR (Qiagen) or fixed at specific developmental stages for further analysis. Morpholinos were purchased from GeneTools. A splice-blocking morpholino (5′-GTAAAATCTCGCTCACATGATCTGC -3′) was selected for knocking down *Rab23*. This splice-blocking MO, which allows the efficiency of MO inhibition to be determined through PCR, was used for all the rab23 morphants shown here. The control morpholino (5′-CCTCTTACCTCAGTTACAATTTATA-3′) used was the scrambled sequence from Gene Tools.

### Whole-mount antibody staining

Whole-mount antibody staining on zebrafish embryos was performed according to standard protocols. Monoclonal anti-acetylated-*α*-tubulin (1:500; Sigma Aldrich) was used to stain cilia. For confocal microscopy, appropriate Alexa-Fluor conjugated secondary antibody was used for signal detection and embryos were counterstained with 4,6-diamidino-2-phenylindole (DAPI) to visualize cell nuclei. Stained embryos were dissected from their yolk and mounted in 70% glycerol. High-resolution images of embryos were captured using a Zeiss LSM 710 confocal microscope (Carl Zeiss Pte Ltd, Singapore).

### Cloning and *in vitro* transcription of capped mRNA

cDNA was synthesized from 24 hours post-fertilization (hpf) wild-type embryos using a cDNA synthesis kit (Invitrogen). The cDNA template was used for PCR amplification of the full-length zebrafish *rab23*. The PCR products were cloned into TOPO vector and subsequently sub-cloned into pCS2-GFPxlt for the rescue experiment. Plasmid encoding *rab23* was linearized, and capped, full-length mRNA was transcribed from this template using the mMessage mMachine Kit (Ambion). The mRNA was injected into one- to two-cell stage embryos in combination with a morpholino.

### Primary cilia length and volume analysis

For the *in vivo* tissue samples, the measurements of cilia volume and cilia length were performed on 3D maximum intensity projection of z-stacked confocal images (at 15-20 µm thickness of z-layers captured at 1 µm intervals) taken from 2-3 spatially matched tissue slices from each embryo, with 3-4 biological replicates in each genotype. For the *in vitro* samples, the measurement of cilia volume and cilia length was performed on 3D projected confocal images (at 10-15 µm thickness of z-layers captured at 1 µm intervals) taken from 2-3 random views in each group. The box plots illustrate data from 3 - 5 independent experiments. The analyses of cilia length and volume were performed on 3D projected images using the ImageJ plugin CiliaQ (Hansen et al., 2021). The default parameters for 3D CANNY-threshold for ciliary reconstruction was applied. Background signal subtraction was applied to animal tissue slices and human NPCs images prior to analysis. In the measurement of cilia length, the data points at below 0.2 μm and above 8 μm were excluded as potential false signals. All data analysis was conducted using at least 3 biological replicates in each genotype.

### Statistical analysis

Unpaired 2-tailed Student’s t-test was performed for statistical comparison between two groups. For comparison of more than two groups, one-way ANOVA, Bonferroni’s Multiple Comparison Test was used. Error bars depict SEM (standard error of the mean). P value: **** *p* ≤ 0.0001, *** *p* ≤ 0.001, ** *p* ≤ 0.01, * *p* ≤ 0.05. Graphs with boxplots depict the interquartile range of the ciliary length or volume, with the lower quartile (Q1) representing the 25th percentile of the data, and the upper quartile (Q3) representing the 75th percentile of the data. The line inside the box indicates the median of the data. For *in vivo* tissue sections analyses on Z-projected confocal images, 3 to 4 confocal images spanning the upper and lower Z-planes of the comparable region of interest were captured with the same imaging parameters, and analysed to represent the result for each animal. In general, 3 to 4 animals were analysed for each genotype, unless stated otherwise. For cell culture experiments, all statistical results were collected from at least 3 to 5 independent cultures.

## Supplemental Information

**Supplementary Figure S1.**
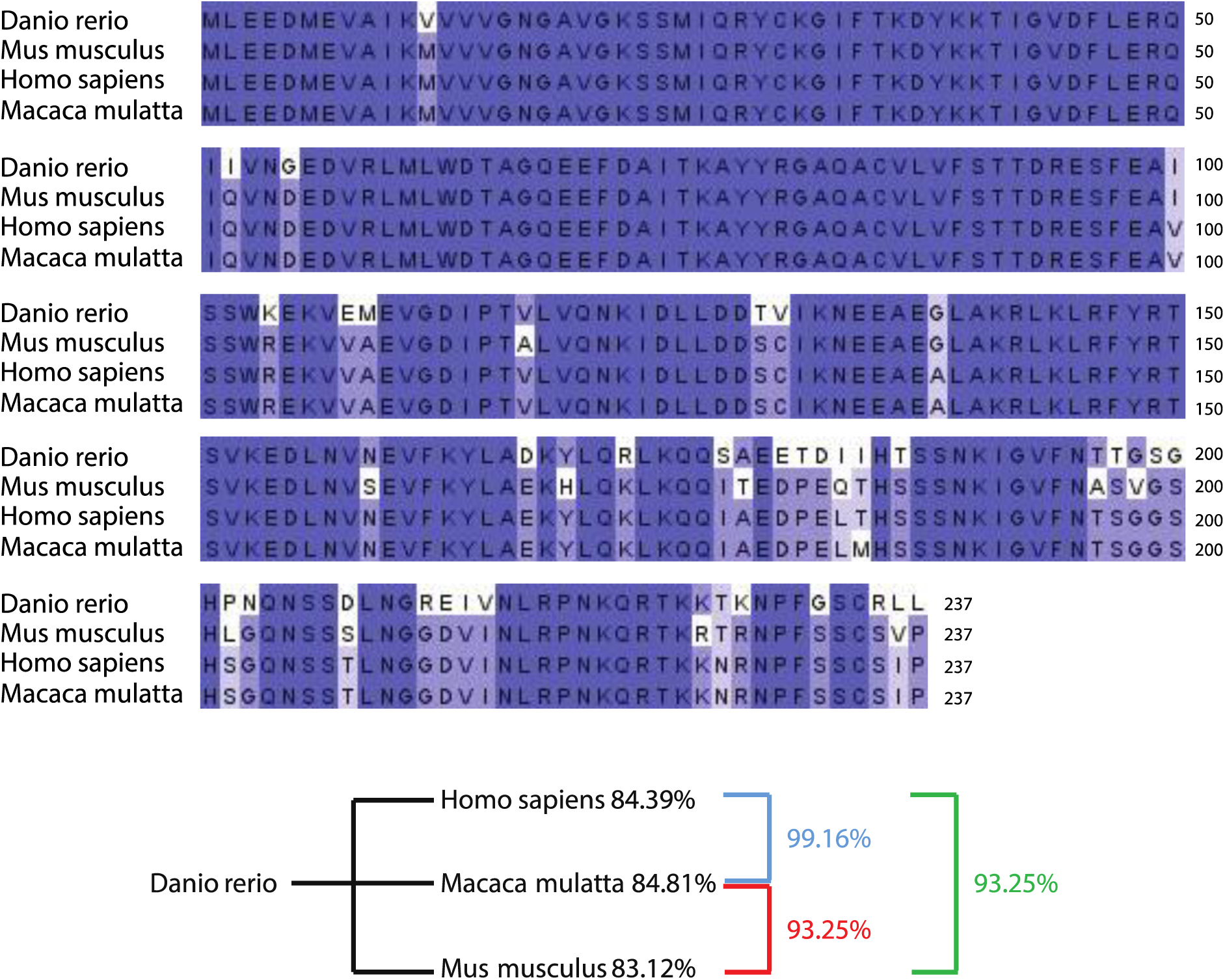
Amino acid sequence alignment reveals highly conserved RAB23 protein across different vertebrates. Multiple protein sequence alignment of RAB23 in zebrafish (*Danio rerio*), mouse (*Mus musculus*), rhesus macaque (*Macaca mulatta*) and humans (*Homo sapiens*). The sequences highlighted in dark blue depict the identical amino acid regions. *Danio rerio* shares 84.39% identity with human RAB23. *Mus musculus* shares 93.25% identity with human.

## Acknowledgement

We thank Bor Luen Tang from the National University of Singapore for the constructive discussions and comments on this work. We thank Eyleen LK Goh for supporting some of the experimental reagents. We thank Wee Lin Wong and Choo Wai Lo for the assistance on figure editing and some data analysis. This study was supported by Research Grant Council-Collaborative Research Fund (CRF-C2103-20GF), HKBU Seed Fund, and the National Medical Research Council–Young Individual Research Grant (NMRC/OFYIRG/0079/2018) to C.H.H. Hor. Work on CS in AOMW’s laboratory was supported by the UKRI Medical Research Council.

## Declaration of Interests

The authors declare no conflict of interest.

## Authors Contributions

C.H.H. Hor conceptualised and conceived the project, wrote the manuscript, performed the human iPSC, human NPCs, animal experiments, immunostainings, confocal imaging, and data analysis. W.Y. Leong performed the zebrafish experiments, amino acid sequence alignment analysis, confocal imaging, data analysis, and wrote the manuscript. W.L. Tung performed the mouse NPC culture, qPCR, immunohistochemistry staining, confocal imaging and cilia analysis. W.H. Chui performed part of the immunohistochemistry staining, confocal imaging and partial cilia analysis of embryonic samples. Andrew O.M. Wilkie provided the clinical samples and edited the manuscript.

## Notes

### Competing Interest Statement

The authors have declared no competing interest.

